# Electrophysiology and transcriptomics reveal two photoreceptor classes and complex visual integration in *Hirudo verbana*

**DOI:** 10.1101/552018

**Authors:** Annette Stowasser, Aaron Stahl, Joshua B. Benoit, Daniel A. Wagenaar

**Author notes:** Correspondence: 1200 E. California Blvd, m/c 139-74; Pasadena CA 91125.

## Abstract

Among animals with complex visual processing mechanisms, the leech *Hirudo verbana* is a rare example in which all neurons can be identified. However, little is known about its visual system, which is composed of several pigmented head eyes and photosensitive non-pigmented sensilla that are distributed across its entire body. Although several interneurons are known to respond to visual stimuli, their response properties are poorly understood. Among these, the S cell system is especially intriguing: It is multimodal, spans the entire body of the leech, and is thought to be involved in complex sensory integration. To improve our understanding of the role of this system, we tested its spectral sensitivity, spatial integration, and adaptation properties. The response of the S cell system to visual stimuli was found to be strongly dependent on the size of the area stimulated, and adaptation was local. Furthermore, a “bleaching experiment” demonstrated that at least two color channels contributed to the response, and that their contribution was dependent on the adaptation to the background. The existence of at least two color channels was further supported by transcriptomic evidence, which indicated the existence of at least two distinct groups of putative opsins for leeches. Taken together, our results show that the S cell system has highly sophisticated response properties, and could be involved in the processing of complex visual stimuli. We propose the leech as a novel system to understand visual processing mechanisms with many practical advantages.

**Summary statement:** Detailed quantitative analysis of light responses in the medicinal leech *Hirudo verbana* unequivocally demonstrates the existence of parallel visual pathways processing visual and UV stimuli. Responses to spatially complex stimuli indicate relatively sophisticated information processing.

## Introduction

Vision requires complex integration mechanisms. In most model species, investigating those at the level of individual neurons is complicated by the large number of neurons involved and the challenge of identifying specific neurons. Among animals with complex visual processing, the leech *Hirudo verbana* is a rare example in which all neurons can be readily identified. However, little is known about the neuronal mechanisms of visual processing in the leech. At the input level, the leech’s visual system consists of several pigmented cylindrical eye cups within the head region, and a grid of nonpigmented photosensitive sensilla distributed across the entire body (Kretz et al., 1976). Several interneurons have been found to respond to visual stimuli (Kretz et al., 1976), but their response properties remain poorly understood. Among these, the S cell interneuron is especially intriguing. The S cell is an interneuron that is activated by salient stimuli of multiple modalities, including mechanical as well as visual stimuli (Magni and Pellegrino, 1978; Laverack, 1969; Bagnoli et al., 1973; Kretz et al., 1976), suggesting that it may be involved in multimodal sensory integration (Harley et al., 2011, 2013). A single (not bilateral) S cell is present in each of the 21 segments of the leech. Synaptic pathways between the S cell and both sensory and motor neurons have been reported within the segmental ganglia (Sahley et al., 1994). Importantly, S cells in adjacent ganglia are strongly coupled by electrical synapses (Frank et al., 1975). The electrical coupling between S cells is so strong that the whole S cell system can be considered as a single syncytium that acts as a fast conducting pathway connecting the segmental ganglia (Peterson, 1984). Although direct proof is lacking (see Sahley et al., 1994), the general consensus in the field is that the S cell system plays a key role in synchronizing general arousal throughout the nervous system of the leech.

Despite the S cell’s purported central role in sensory processing, the neuronal pathways leading from photoreceptor cells to the S cell are not known. In addition, other basic questions regarding the S cell system, including its role in light adaptation, its temporal and spatial integration properties, and its overall role in vision remain to be addressed.

It has long been known that *Hirudo* has the ability to visually detect the direction of water waves, and that—in combination with mechanical cues—it uses this information for prey localization (Dickinson and Lent, 1984; Carlton and McVean, 1993; Harley et al., 2011). This demonstrates that its visual system has the ability to process spatiotemporal patterned visual stimuli, despite the lack of image-forming eyes. S cells respond when the leech is presented with flashes of light as well as to complex stimuli associated with water waves (Lehmkuhl et al., 2018). Their multimodal response properties, along with the fact that the S cell system spans the entire body, makes them an intriguing candidate for further investigations.

Not much is known about temporal properties of S cell responses. However, an early study (Laverack, 1969) found that the S cell response fairly quickly ceases when the leech is stimulated continuously either visually or mechanically, but that the S cell remains sensitive to tactile stimuli when visually desensitized. This finding is consistent with findings in other animals. For instance, the human central nervous system is well known to fairly quickly adapt to constant or repetitive stimuli, while remaining sensitive to different stimuli of the same or a different modality. This phenomenon is generally believed to enhance an animal’s ability to detect ethologically relevant changes in its environment (Desimone and Duncan, 1995), though much about the underlying mechanisms remains to be fully understood.

Another early study of leech vision (Kretz et al., 1976) indicated that a single class of photoreceptors is involved in responding to light. Those photoreceptors, putatively found in both the head eyes and the sensilla, respond most strongly to light in the green range of the visual spectrum. Unexpectedly, however, recent behavioral and electrophysiological experiments demonstrated that under certain specific circumstances, the S cell system responded more strongly to UV than to green light. This phenomenon was observed especially when UV light was directed at the ventral side of the body wall, suggesting that the S cell system may play a role in detecting and correcting the animal’s orientation relative to the sun (Jellies, 2014a, b).

These results appear to require the presence of a second class of photoreceptors, which have not been directly identified. There is, however, precedence for the existence of multiple photoreceptor classes in other leeches: molecular investigations in *Helobdella robusta* have found at least four distinct opsins (Döring et al., 2013). Unfortunately, the spectral properties of these opsins remain unknown due to a lack of physiological and molecular data.

In this paper we present electrophysiological and transcriptomic evidence indicating the presence of at least two distinct photoreceptor classes in *Hirudo*. Furthermore, we show that the S cells are involved in spatial integration and the implementation of differential adaptation to background light illumination, unveiling new roles for the S cell system in vision and sensory integration.

## Materials and methods

### Animals and animal preparation

Adult leeches (*Hirudo verbana*) were obtained from Niagara Leeches (Niagara Falls, NY, USA) and maintained under standard conditions (Harley et al., 2011). At the time of experiments, leeches had fasted for at least two months and weighed 1–1.5 grams. Leeches were anesthetized with ice cold leech saline (Tomina and Wagenaar, 2018) and immobilized ventral-side up on dark silicone (Sylgard 170, Dow Corning, Midland, MI, USA) using insect pins stuck in annuli without sensilla. The body wall was opened from mid-body segments from M8 to M11 (or from M7 to M10 in experiments on spectral sensitivity under full dark adaptation). The lateral roots of ganglia M9 and M10 (or M8 and M9) were transected, and the ganglia and connectives were gently separated from the body tissue without severing any other nerves. The wall of the ventral blood sinus (“stocking”) was removed between the exposed ganglia. A thin strip of silicone (Sylgard 184) was slipped between the nerve cord and the body wall and pinned down on each side of the leech. Ganglia M9 and M10 (or M8 and M9) were pinned very close together onto the silicone strip and the connective between them was sucked into a suction electrode. The general setup is shown in Fig. S1A. The temperature of the leech was kept at 15–19 °C throughout all experiments.

### General Electrophysiological Setup

The electrophysiological setup consisted of a differential amplifier (Model 1700, A-M Systems, Sequim, WA, USA), an oscilloscope (TBAS 1046, Tektronix, Beaverton, OR, USA), and an A/D converter (Model 118, iWorks Systems, Dover, NH, USA). Recordings were performed inside a Faraday cage on a vibration isolation table (TMC 66-501, Technical Manufacturing Corporation, Peabody, MA, USA). Data was stored on a PC using LabScribe software (iWorks), and analyzed using custom-written code in Octave (Eaton et al., 2017). To tightly control background illumination, the entire recording area was enclosed in black-out fabric (BK5, Thorlabs, Newton, NJ, USA). In addition, the room light was kept off during experiments, so that the only light sources in the room where indicator lights on electronic equipment and a computer screen. The light seal of the recording area was tested by means of a sudden substantial increase in ambient room light after the leech was fully dark adapted and verifying that this did not elicit a response.

### Measuring Light Intensities

Measurements were taken with a spectrometer (USB2000+ with a QP600-025-SR optical fiber and a CC3-UV-T cosine corrector; Ocean Optics, Dunedin, FL, USA) which was calibrated against a calibrated light source (DH-2000, Ocean Optics). All reported light intensities are absolute numbers from radiometric irradiance measurements, in units of photons/cm^2^/s. To obtain controlled light intensities below the minimum intensity that the spectrometer could directly measure, we used calibrated neutral density filters placed in front of a brighter light source. Calibration of neutral density filters was performed independently for each relevant wavelength. All measurements were made with the cosine corrector of the spectrometer probe at the same distance and orientation relative to the light source as the leech would be in our actual experiments. Although we took great care to measure light intensities as accurately as possible, it should be noted that measuring absolute light intensities accurately is notoriously challenging: according to Johnsen (2012), measurement errors up to 10% (0.1 log units) are to be expected even in the best of scenarios. We believe our measurements to be accurate to about that level. Furthermore, since all of our key results rely on relative light intensities, minor errors in absolute intensity values do not affect the interpretation of our results.

### Spectral sensitivity measurements

Monochromatic light was generated by coupling a 150W Xenon arc lamp (Apex 70525 Monochromator Illuminator, Oriel Instruments, Stratford, CT, USA) to a monochromator (Cornerstone 130 1/8m 74000, Oriel). In previous experiments using this system, we had observed a small secondary peak at approximately 300 nm below the primary peak wavelength. To eliminate this secondary peak, we used a long-pass filter (ET542LP, Chroma, Bellows Falls, Vermont, USA) for all primary wavelengths of 590 nm and above. The light intensity was controlled with a variable neutral density filter (50Q00AV.2, Newport Corporation, Irvine, CA, USA) mounted on a motorized rotator stage (NSR-12 controlled by a NewStep NSC200 controller, both Newport). Three additional neutral density filters (FRQ-ND1 and FRQ-ND2, Newport; NDUV30A, Thorlabs) that were mounted onto a manual filter wheel (FW1A, Thorlabs) were used to achieve light attenuation beyond the range of the motorized filter wheel. The duration of the stimulus was controlled with a shutter (VCM-D1, Uniblitz, Rochester, NY, USA). The light path also contained two lenses (LJ4395-UV and LA4306-UV, Thorlabs) that focused the light onto an optical fiber placed directly behind the shutter. (Lenses and fiber were chosen to transmit both UV and visible light.) At the end of the optical fiber was a lens that collimated the light so that a light spot with a diameter of 2.8–3.5 cm was projected onto the leech from a distance of 10–13 cm. The light source was positioned above the leech and illuminated the entire posterior ventral side ranging from the body wall opening at M10 to the rear sucker at an angle of no greater than 30° from normal.

Leeches were dark adapted for at least 30 minutes before starting a recording, and recordings were performed without background illumination. (We could not quantify stray background light, but estimate it to be below 10^8^ photons/cm^2^/s, or approximately 0.0002 lux, similar to the darkness under an overcast sky on a moonless night). We recorded responses to 500-ms stimuli with the following peak wavelengths (in nm): 320, 350, 400, 455, 530, 590, and 655. The order of wavelengths that we tested was randomized. To generate response–log(intensity) curves, we used light intensities in a range of approximately 3 log units in steps of approximately 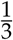 log units, always working in order of increasing light intensity, separately for each wavelength. Preliminary data (not shown) showed that it was critical to leave prolonged recovery times between stimuli especially after a strong response to relatively high light intensity. To optimize for quality of obtained data, we allowed at least 1 minute and up to 5 minutes between stimuli, depending on the stimulus light intensity and responses.

### Adaptation to green and UV

For these experiments, we used LEDs in combination with neutral density filters to achieve higher light intensities and a wider range of intensities than what was possible with the monochromator. The LEDs were controlled by a custom driver that provided a precisely regulated DC current to the LED; the neutral density filters served to extend the intensity range beyond the range of the driver. We specifically did not use pulse width modulation (i.e., control of the duty cycle of flicker) to avoid assumptions about the frequency response of the visual system. Schematics are available on request.

For UV, we used LEDs with a dominant wavelength of 365 nm (LED Engin LZ1-10UV00, Mouser) and the same ND filters as before. For green light, we used 523-nm LEDs and OD-2 and OD-4 filters (NE20B-A and NE40B-A, Thorlabs). In this way, we achieved a green background light intensity range of 6 log units and a green and UV stimulus light intensity range of approximately 7.5 log units. The UV stimulus light (but not the UV background) was fitted with a filter (357/25x, Chroma AT) that eliminated a small secondary peak within the visual wavelength range. Since UV illumination elicited a strong fluorescence of the exposed intestinal tissue at the body wall opening, we removed this tissue as best as possible, and closed the body wall opening up as much as possible for the recording.

Each LED was mounted behind a condenser lens (ACL2520, *f* = 20 mm, Thorlabs). The background and stimulus LED assemblies were mounted directly above the leech such that the angle between them was no more than 15°. The background illuminated the leech from a distance of 19 cm; the stimulus from a distance of 11 cm. The illuminated area had a diameter of 9.5–10.5 cm. The leech was pinned out to a length of no more than 6 cm, so that the entire ventral side of the leech was illuminated by both the background and the stimulus (Fig. S1B). The green and UV background LEDs were mounted at fixed positions immediately adjacent to one another on a slider that allowed their positions to be switched. This ensured that the stimulus location and orientation was identical regardless of wavelength.

To quantify the adaptation to green background light, we tested six background intensities ranging from 3.4 ×10^10^ to 3.4 ×10^15^ photons/cm^2^/s in steps of one log unit. Because the need to keep our experimental animals healthy throughout the experiment imposed time constraints on the duration of experiments, each leech (*N* = 11) was tested with only three or four of the six background light intensities. (Specifically, we tested the lowest light level on 11 leeches, the second level on 6 leeches, the third on 4, fourth on 3, fifth on 5, and highest on 10.)

As before, leeches were dark adapted for 30 minutes before recording, and additionally background adapted for 10 minutes every time we changed the background illumination or had to open the light seal to exchange neutral density filters. To generate response–log(intensity) curves for each background light intensity and stimulus wavelength (green and UV), we applied 2-second stimuli with intensities spanning 3 log units in steps of approximately 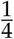 log units, in order of increasing light intensity. To prevent adaptation to the stimulus intensity, 3 minutes of only background illumination was provided between stimuli.

### Local versus non-local adaptation

Two green through-hole LEDs (941-C505BGANCC0D0781, Mouser) provided differential background illumination to the anterior and posterior halves of the leech. A third such LED delivered flash stimuli to the posterior half of the leech. All LEDs were mounted at a distance of 9 cm from the leech; the stimulus LED was mounted immediately adjacent to the LED that provided background illumination to the posterior half of the animal. A light barrier consisting of blackout fabric was placed between the anterior and posterior halves of the leech to ensure controlled differential stimulation of the two halves (Fig. S1C). As before, we used neutral density filters to reduce light intensity beyond the range of the LEDs. These were mounted onto a slider so that they could be exchanged from the outside without opening the light seal of the recording area.

Two levels of background light intensity were used in these experiments: 3.9 ×10^12^ photons/cm^2^/s (“dark”) and 4.4 ×10^13^ photons/cm^2^/s (“light”). All combinations of light and dark background conditions were tested, always in the following order: 1. Both halves dark; 2. Both halves light; 3. Posterior light, anterior dark; 4. Posterior dark, anterior light; 5. Both halves dark (as a control to test if we could recover the initial response). For constructing response curves, the same range, step size, order of stimulation, and stimulus duration was used as for the previous experiment.

### Spatial integration

Background illumination intensity was 4.5 ×10^11^ photons/cm^2^/s. The setup was otherwise the same as for the local versus non-local adaptation experiment, except that an additional stimulus LED was used to provide flashes to the anterior region. Order of stimulation was: 1. Anterior only; 2. Anterior and posterior together; 3. Anterior only again to test if we could recover the initial responses. After that we cut the cord posterior to the recording site, which disconnected the posterior half of the body from our recording site, and tested for the influence of stray light by stimulating: 4. Posterior only (which potentially could affect the anterior side through stray light); 5. Anterior only, to test if initial responses could be recovered. Stimulus duration, intensity range, step size, order of stimulation and time between stimuli were as before.

### Data analysis

Action potentials from the S cell were identified as the largest spiking units in extra-cellular recordings from the nerve cord (Frank et al., 1975). Electrophysiological data were analysed using custom programs in Octave. As a measure of response strength, we counted S cell spikes that occured within a certain time window, starting when the stimulus was turned on. This time window was either as long as the stimulus duration (spatial integration and local versus non-local adaptation experiment), or slightly longer (spectral sensitivity experiment: 1.5 s; adaptation to background experiments: 2.5 s). Response– log(intensity) curves are standard logistics:

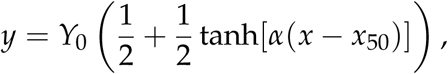

where *y* is spike count, *x* is log intensity, *Y*_0_ is the maximal spike count (plateau response), *α* is the slope of the curve, and *x*_50_ = log(*I*_50_) is the light intensity (in log units) that elicits half maximal response. For quantifying the light intensity for 50% response (*I*_50_, Figure 1), the plateau spike count (*Y*_0_) was determined once per leech and then used for all wavelengths. Likewise in Figure 2, the plateau spike count was determined once per leech (for green stimuli) and used for all background levels and both UV and green stimuli. The same principle was used subsequent figures, except that in Figure 3 we used 35% of maximum as the critical value, because UV-on-UV stimuli often did not elicit 50% of maximum green-on-green response even at the highest intensities. To find the delay of the response (Figure 2), we measured the time that elapsed from the beginning of the stimulus to the occurence of the 3rd spike of the response.

**Figure 1.**
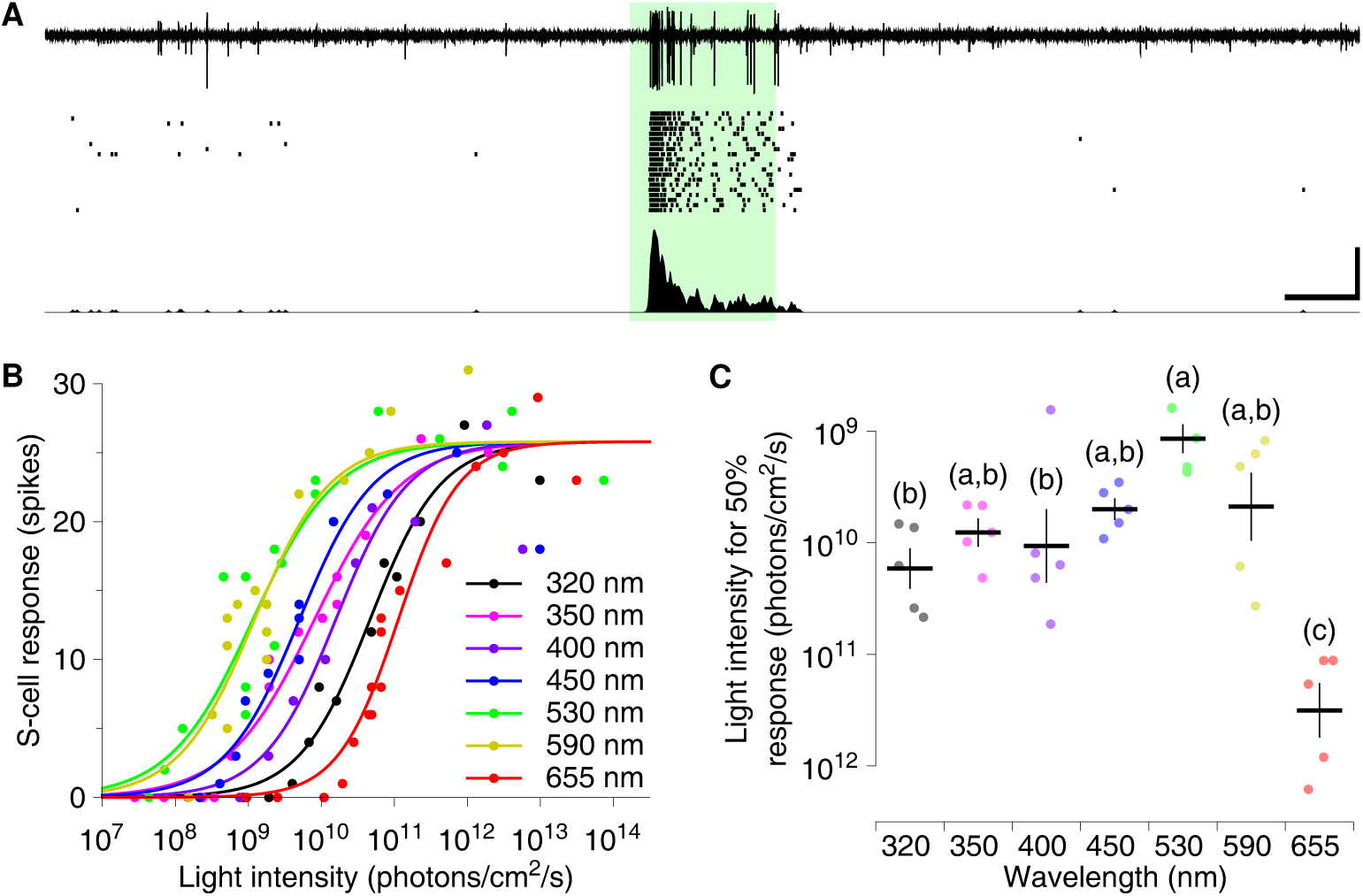
S cell responses to light stimulation and spectral sensitivity. **A.** Responses to 2-s-long flashes of green light (530 nm, 1.5 ×10^11^ photons/cm^2^/s) presented to the posterior half of the ventral body wall. *Top to bottom:* representative raw extracellular trace; raster plots from 20 individual trials on a single leech; firing rate histogram of those trials. Scale bars: 1 s and 25 spikes/s. **B.** Response curves to 500-ms-long flashes of light of various wavelengths (one representative leech). **C.** Spectral sensitivity of S cell responses. Dots represent individual animals; black lines mark means and standard errors. Letters mark groupings from ANOVA/Tukey (at *p* < 0.05; *n* = 5).

**Figure 2.**
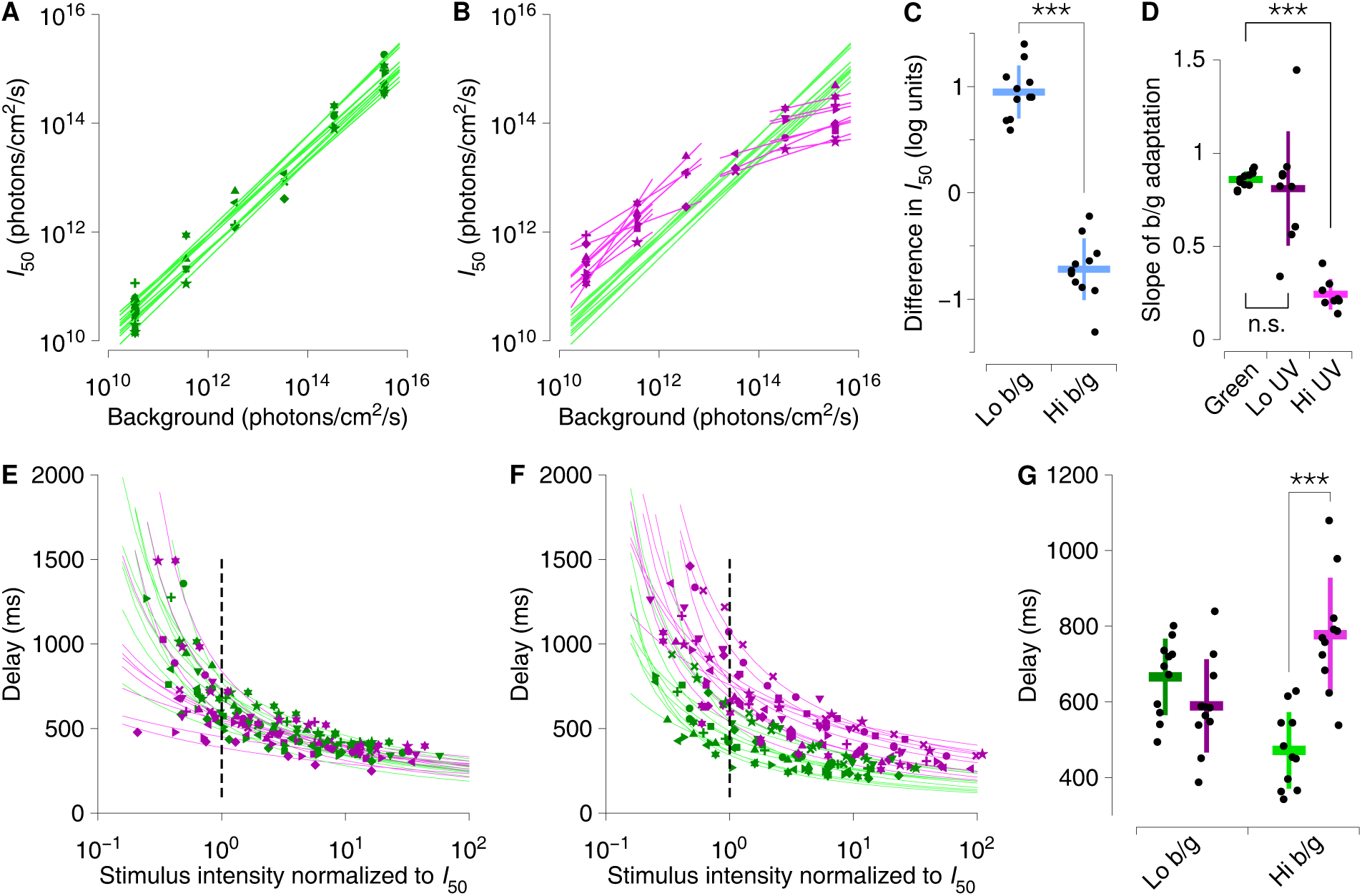
Adaptation to green background light. **A.** Intensity of green stimulus light required to attain 50% of plateau response (*I*_50_, see text) as a function of green background intensity. Symbols indicate animals; lines are linear fits for each animal. **B.** Intensity of UV stimulus light required to attain the same response as in (A) as a function of green background intensity. Lines are linear fits separately for the low-background and high-background regimes. **C.** Difference in light intensity required to attain 50% of plateau response using UV light vs. green light, at the lowest background intensity (*left*) and at the highest background intensity (*right*). ***: *p* < 10^−7^, t-test (*n* = 11). **D.** Summary of fit results from (A) and (B): Black dots indicate the slopes of individual fits; bars indicate mean and standard deviation across animals. ***: *p* < 10^−5^, t-test (*n* = 8). **E & F.** Delay of the response to green and UV light stimulation at lowest background light intensity (E) and at highest background light intensity (F). The stimulus light intensity is plotted normalized to *I*_50_. Symbols indicate individual leeches, lines are fits for each animal, the broken line indicates the light intensity that elicited half-maximum response (*I*_50_). **G.** Summary of the data shown in (E) and (F), showing the delay of the response at *I*_50_ for green and UV stimulation at lowest and highest background light intensity. **: *p* < 0.005, ***: *p* < 10^−5^, t-test (*n* = 11).

**Figure 3.**
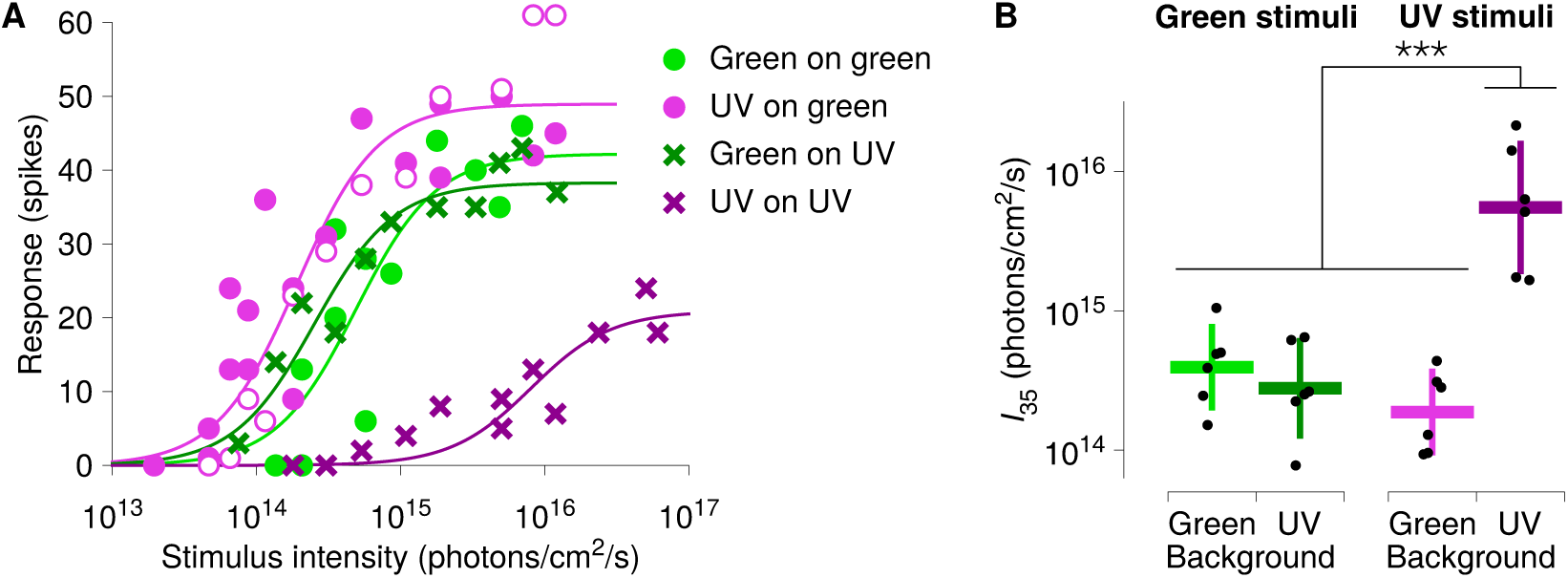
Adaptation to the spectrum of background light. **A.** Response to stimuli with green light (shades of green) and UV light (shades of pink and purple) on a background of either green light (disks) or UV light (cross marks). Background intensities were 3.4 ×10^15^ photons/cm^2^/s for the green background, and 9.7 ×10^15^ photons/cm^2^/s for the UV background (see text for rationale). Closed and open pink markers represent data collected under the same conditions at the beginning and end of the experiment, to confirm stability of responses. Data from one representative animal. **B.** Stimulus light intensity required to elicit a response at least 35% as strong as the plateau response for UV stimuli on green background. ***: *p* < 0.0001, Tukey test following ANOVA (*F*_3,15_ = 32.5, *p* < 10^−6^, *n* = 6 leeches). Colors as in (A).

### Transcriptome analyses to identify opsins

Transcriptomic databases were generated from two separate tissue types: a single head containing the eyes, and 100 isolated sensilla collected from the body. Tissues were dissected in ice-cold RNAse free Gibco PBS (Thermo Fisher Scientific, Waltham, MA, USA). Tissues were briefly frozen in liquid nitrogen and ground using a mortar and pestle. RNA isolation was conducted using the RNeasy Lipid Tissue Kit (Qiagen, Valencia, CA, USA). To assess the quality of RNA, extractions were subjected to spectrophotometric analysis utilizing a NanoDrop 1000 Spectrometer (Thermo Fisher Scientific, MA, USA) where the A260/280 absorbance ratio yielded measurements around 2.0 for RNA extracts, indicating that all RNA measurements were relatively pure. RNA-seq utilized the Illumina HiSeq 2500 (75 bp) with Ribo-zero preparation at Cincinnati Children’s Hospital Core Sequencing Facility (Cincinnati, OH, USA). The raw read FASTQ files were assembled through the utilization of Trinity Grabherr et al. (2011), CLC Genomics, and Oases (Schulz et al., 2012) according to previously described methods (Rosendale et al., 2016). Expression was assessed by mapping reads based on parameters described in Rosendale et al. (2016). Quality of each transcriptome was assessed through evaluation of the Benchmarking Universal Single-Copy Orthologs (BUSCO) gene sets (Simão et al., 2015).

Opsin sequences were identified using the Blastx algorithm (Altschul et al., 1997) to identify orthologs to the previously annotated opsin sequences of *Helobdella robusta* (Döring et al., 2013) along with opsin sets obtained from arthropod and other invertebrate species from NCBI nr databases. These two different databases were used to identify potential functionality, as many annelid-specific opsin have not been fully characterized. A reciprocal BLAST against the invertebrate and arthropod databases was used to confirm if predicted genes match opsins in other systems. Relationships between the opsin sequences and contigs was assessed through the use of MEGA5 Tamura et al. (2011) to generate a neighbor joining tree after sequence alignment with CLUSTAL Omega (Sievers and Higgins, 2014). Illumina sequencing files have been deposited to the NCBI SRA (Bioproject: PRJNA504032).

## Results

To examine the light dependent responses of the S cells, we investigated their response strength as a function of the wavelength of the stimulus and adaptation to background illumination, tested whether the adaptation to background illumination is local or global, and quantified spatial integration. We focused specifically on the S cell’s response to light stimulation of the ventral body wall.

### General response properties

The S cell system responded reliably and vigorously to stimulation of the ventral body wall of the leech with flashes of light. The typical response to a flash of green light presented against a dark background is illustrated in Figure 1A. The response can be separated into two phases: a) an initial transient phase characterized by high firing rates and b) a sustained response that typically lasts beyond the duration of the stimulation with a substantially lower spike frequency.

### Spectral sensitivity of dark-adapted leeches

To test the spectral sensitivity of the S cell system, we applied 500-ms flashes of light of various wavelengths and intensities to the ventral body wall of dark-adapted leeches and recorded spike responses from the S cell using suction electrodes. For each wavelength, we constructed response–log(intensity) curves by fitting logistic functions (Figure 1B). We then quantified the light intensity required to elicit half the maximum response for each leech to obtain absolute sensitivity profiles (Figure 1C). In agreement with Kretz et al. (1976), we found the highest sensitivity in the green wavelength range. We also observed a small secondary peak in the UV range (around 350 nm), although the statistics were inconclusive. Certainly, these dark-adapted leeches failed to show the strong response to UV light reported by Jellies (2014a).

### Physiological evidence for a second photoreceptor

To investigate whether the putative secondary peak corresponded to a second photoreceptor type, we performed a series of background adaptation or “bleaching” experiments designed to unmask subtle secondary responses that otherwise remain hidden by the strong response of the green-sensitive photoreceptor. We argued that increasingly intense green background light would increasingly bleach out the green-sensitive photoreceptor, so that increasingly strong flashes would be needed to activate it, regardless of the color of those flashes. In contrast, the effect on a possible second photoreceptor that is only sensitive to UV light would be minimal.

Thus we began by adapting leeches to a variety of background intensities of green light and measuring response curves to flashes of green light superimposed on those backgrounds. We found that over a range of nearly 6 log units, the response was approximately contrast invariant, that is, the intensity for half-maximum response, or *I*_50_, scaled almost in direct proportion to the background intensity (Figure 2A): the slopes of the best fit lines are 0.86 ± 0.04 (mean ± SD, *n* = 11 animals; Figure 2D).

We also presented these leeches with flashes of UV light against the same green backgrounds, and found that at low background intensities (up to 10^12^ photons/cm^2^/s), the intensity required to obtain half-maximum response again scaled nearly proportionally with the background intensity (Figure 2B, left half). The best fit lines had slopes of 0.81 ± 0.31 (mean ± SD; *n* = 9 animals), not significantly different from the “green” slopes (t-test). This indicates that the responses to UV light were due to the same photoreceptor that was bleached out by the green background light.

However, this trend did not continue at higher background intensities: At green background intensities above 10^14^ photons/cm^2^/s, the intensity of UV light required to obtain half-maximum response no longer increased linearly with the background intensity at all (Figure 2B, right half). In this range, the best fit lines had slopes of 0.24 ± 0.08 (*n* = 8), and at the very highest green background intensities, sensitivity to UV flashes was actually greater than to green flashes (Figure 2C), suggesting that a second photoreceptor is activated by high-intensity UV light.

Further evidence for the involvement of two photoreceptors in the S cell response comes from analyzing response delays: At the lowest background light intensity, the delay of the response to UV stimulation was similar to the delay to green stimulation (on average 589 ± 124 vs. 666 ± 101 ms, mean ± SD, Figure 2E and G), whereas at the highest green background light level, the delay of the response to UV was substantially longer than the delay of the response to green stimulation (on average 777 ± 151 vs. 471 ± 101 ms; *t*_10_ = 8.8, *p* < 10^−5^; Figure 2F and G). This difference could easily be explained if the two photoreceptors have distinct temporal response properties or connect to the S cell via two pathways that introduce distinct delays. It would be harder to explain if there were only one photoreceptor type.

We next performed a direct test for the presence of two distinct color channels (viz., UV and green) that contribute to the responses in high-intensity background light: We presented leeches with flashes of green or UV light on top of the highest intensity green background from the previous experiment, and also with those same flashes presented against a bright UV background. We purposefully chose the intensitiy of that UV back-round light such that green flashes against this background elicited similar responses as against the green background (Figure 3A, green curves). As expected, this required more photons of UV than green background light (9.7 ×10^15^ UV photons/cm^2^/s vs 3.4 ×10^15^ green photons/cm^2^/s of green light): this merely confirmed that a substantial contribution to the response to green flashes came (largely) from a receptor that was more sensitive to green than to UV light, and hence was also more susceptible to bleaching by green light than by UV light. Also in agreement with the previous experiment, UV flashes elicited slightly more spikes at slightly lower stimulus intensities against the green background than did green flashes (Figure 3A, pink circles and curves). But crucially, UV flashes elicited far *fewer* spikes against the UV background (purple cross marks and curve), even at very high intensities. This phenomenon was robust across animals: the photon flux required to elicited an equal response using UV flashes against a UV background was significantly larger than when using UV flashes against a green background or when using green flashes against either background color (Figure 3B). The most parsimonious explanation is that the UV background specifically bleached out a (mainly) UV-sensitive photoreceptor.

### Transcriptomic confirmation of a second photoreceptor

To obtain independent confirmation that the observed responses were indeed due to two photoreceptors, we searched transcriptomes for putative opsin genes. We obtained these transcriptomes by performing RNA-seq on a tissue sample from the head (focusing on the head eyes) and on a tissue sample containing 100 sensilla isolated from the body wall. The quality of the resulting transcriptome was evaluated using three BUSCO gene sets (see *Methods*). BUSCO scores were over 80% for all assemblies and above 95% when the three sets were combined (Figure 4A). This indicates that our *de novo* contig library has the completeness required for subsequent analyses.

**Figure 4.**
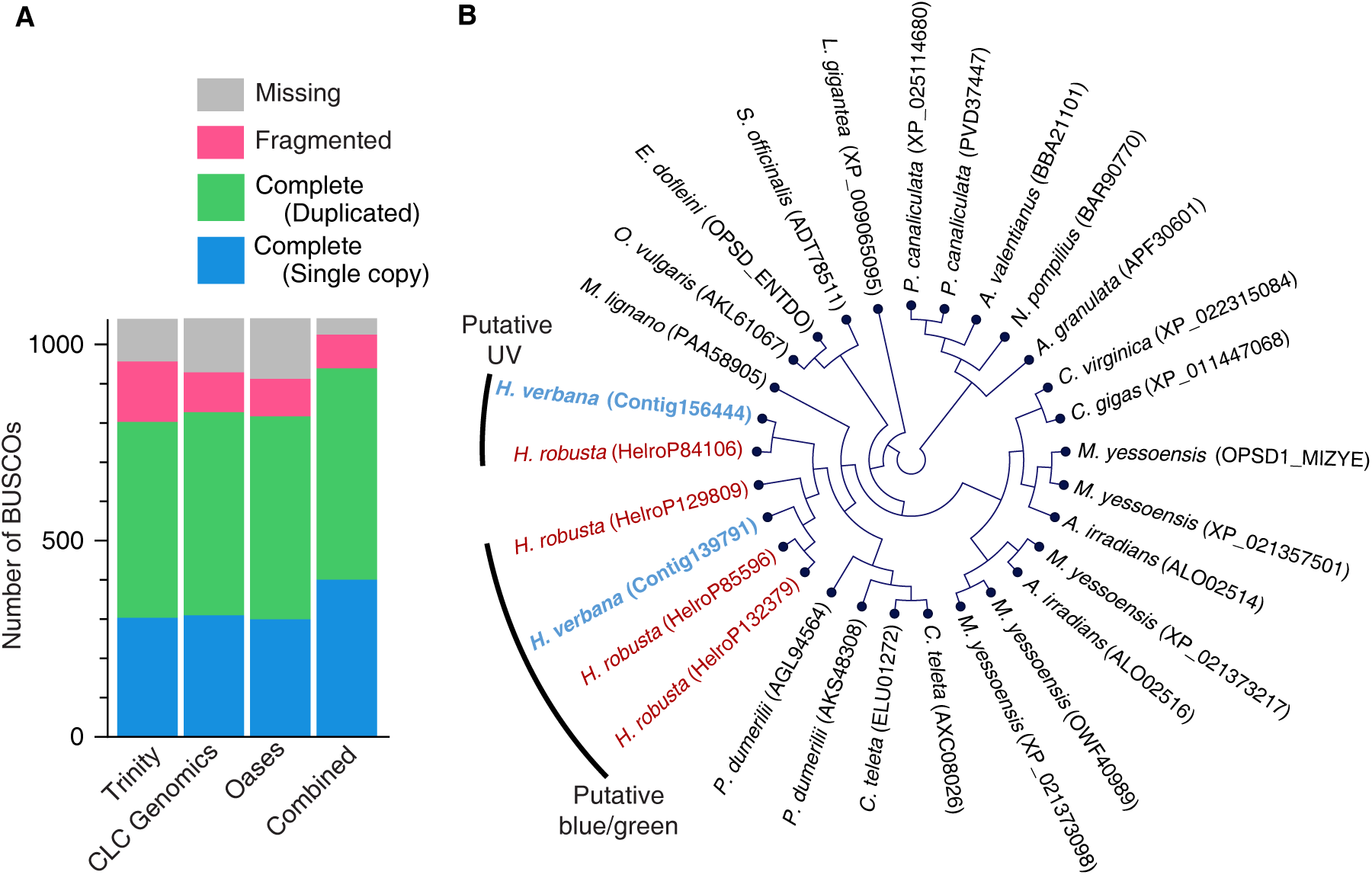
Transcriptome analysis of putative opsins. **A.** BUSCO-based quality assessment of contigs from de novo assembly. **B.** Amino acid phylogeny based on alignment with CLUSTAL followed by sequence analysis and tree generation through the use of MEGA5 (Tamura et al., 2011). All nodes have at least 60% support. Colored names indicate leech opsins.

Two putative opsins from *Hirudo verbana* were identified through BLAST analyses against opins from other invertebrates (Döring et al., 2013), and both had orthologs in another leech (Figure 4B). Each of these had documented expression in both the head and the sensilla. Of the two putative opsins found in *Hirudo*, one (Contig139791, Supplemental File) had BLAST hits with other invertebrate opsins outside of leeches that are sensitive to blue and green wavelengths; the other (Contig156444, Supplemental File) showed similarities to UV opsins from arthropods. We also performed a direct BLAST comparison against a previously described UV-sensitive from another annelid, *Platynereis dumerilii* (Tsukamoto et al., 2017), and found a close match between it and our putative UV opsin (Table S1).

Orthologs of both our putative opsins in *Helobdella robusta* showed similar results: three were likely blue-and green-sensitive and one has putative UV sensitivity. In all, these transcriptomes suggest the presence of one blue-or green-sensitive opsin in *Hirudo* and one UV-sensitive opsin, supporting our physiological experiments.

### Background adaptation affects only local sensory processing

Our experiments thus far showed that S cell responses adapt to background light intensity. However, they did not show whether adaptation occurs in the sensory periphery, in the central nervous system, or in both. In addition, if adaptation occurs in the nervous system, it could be a local phenomenon (limited to the segment or segments targeted by the light), or it could be a global phenomenon (in which illumination of one or several segments would trigger adaptation throughout the animal).

To investigate these scenarios, we differentially adapted the anterior and posterior half of the ventral side of the body wall to two distinct green background light levels and tested the response to posterior green stimulation (*n* = 5). As before, for each animal we established response curves as a function of log intensity (Figure 5A) and calculated the light intensity that elicited 50% of the plateau response (*I*_50_; Figure 5B). As expected, the *I*_50_ for posterior stimulation strongly depended on the background light level on the posterior body wall (blue vs. black points, or red vs. green points; ANOVA, *F*_1;20_ = 55, *p* < 10^−6^). In contrast, the background light level on the posterior body wall had no effect (green vs. black points, or red vs. blue points). Thus, adaptation appears to be a local phenomenon.

**Figure 5.**
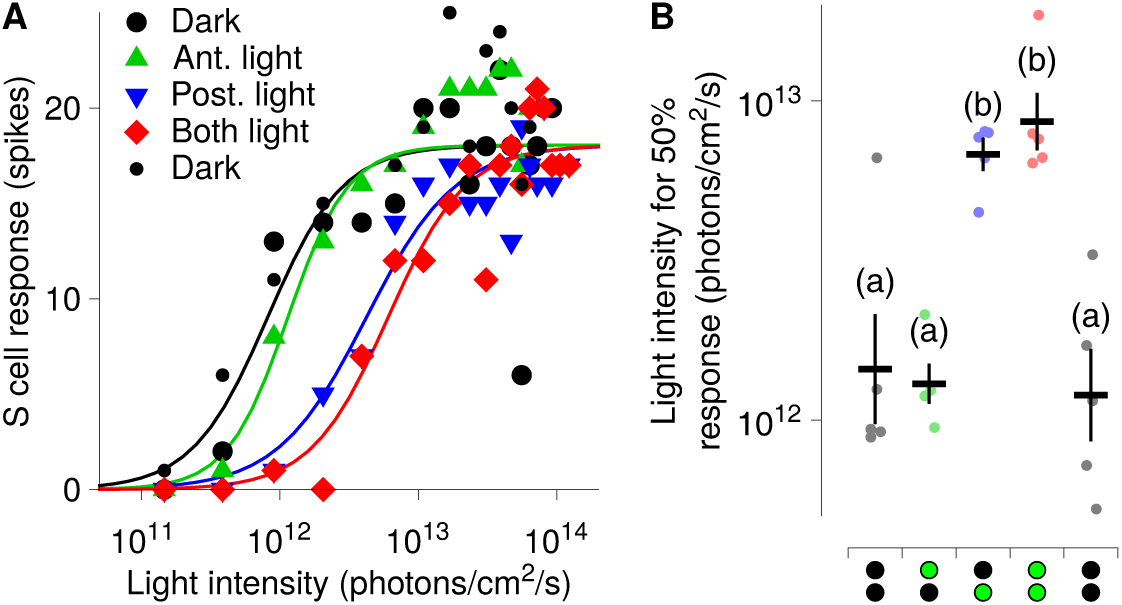
Local and non-local adaptation in the S cell to green background light. **A.** Responses to flashes of light presented to the posterior portion of the ventral body wall when the whole leech was dark-adapted (*black circles*), when the anterior was light-adapted (*green upward triangles*), when the posterior was light-adapted (*blue downward triangles*), and when the whole leech was light-adapted (*red diamonds*). *Small black circles* represent a final repeat of the dark-adapted condition at the end of the experiment to confirm stability of responses. **B.** Intensity of light required to obtain 50% of the maximum response to posterior stimulation, under the same series of conditions used in (A). Symbols below graph serve as mnemonics for light (*green*) and dark (*black*) adaptation for anterior (*top symbol*) and posterior (*bottom symbol*). *Dots* represent individual animals (*N* = 5); *black lines* mark means and standard errors.

### S cell responses integrate spatial information

The absence of nonlocal adaptation does not rule out the possibility that the S cell system performs spatial integration. In fact, the intersegmental connections between S cells uniquely position the S cell system to integrate information across the whole nervous system. To investigate that possibility, we stimulated either the whole leech or only the anterior half of the leech with green light while recording from an S cell located in the anterior half. We found that stimulating both halves together elicited a stronger response (Figure 6). This indicates that the S cell system integrates information pertaining to light stimuli from across the body.

**Figure 6.**
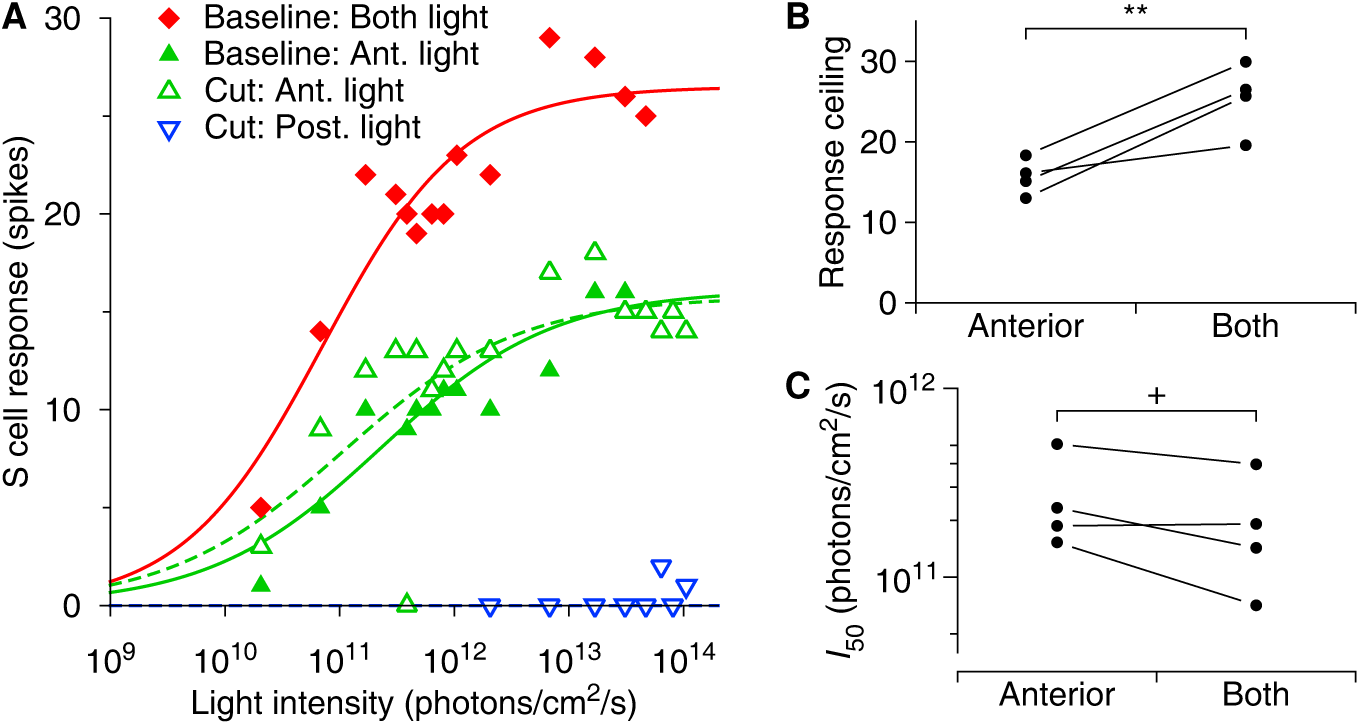
Spatial integration in the S cell. **A.** Response to light flashes presented to the posterior portion of the ventral body wall (*solid blue downward triangles*), anterior portion (*green upward triangles*), or both simultaneously (*black diamonds*). The posterior portion was tested first (*dark blue*) and again last (*pale blue*). Open symbols indicate responses after transecting the nerve cord immediately anterior to the recording site: posterior area stimulated (*red*) or anterior area stimulated (*blue*). Background light level was always 4.5 ×10^11^ photons/cm^2^/s. **B.** Maximum response (plateau of fitted curve) to light flashes presented to anterior and posterior portions simultaneously was higher than to flashes presented only to the anterior portion of the ventral body wall (*t*_3_ = 4.6, *p* < 0.01, one-sided test; *N* = 4). **C.** Light intensity needed to elicit 50% of the respective maximum responses for stimuli presented to anterior and posterior portions simultaneously tended to be lower than for stimuli presented to the anterior portion of the ventral body wall only (*t*_3_ = 2.2, *p* = 0.06, one-sided test; *N* = 4).

To confirm that this integration occurs in the nervous system and that the responses are not merely due to stray light from the posterior illumination reaching sensilla in the anterior part of the animal, we performed control experiments in which we transected the nerve cord posterior to the recording site. Transecting the cord had no significant effect on responses to anterior stimulation, whereas posterior stimulation after transection was completely ineffective (except at extremely high light levels), confirming that the integration is indeed internal (Figure 6A, open symbols).

To quantify these observations, we established response–log(intensity) curves as before. These curves indicated mainly a difference in the plateau (max) response (Figure 6B) with a significantly stronger response to stimulating the whole leech vs only the anterior half (*t*_3_ = 4.6, *p* < 0.01, one-sided test, *n* = 4). The light intensity needed to elicit 50% of the respective plateau responses tended to be marginally lower when the whole animal was stimulated (Figure 6C, *t*_3_ = 2.2, *p* = 0.06, one-sided test, *n* = 4).

In one animal (data not shown) we additionally stimulated the posterior half by itself before transection, which elicited a strong response.

## Discussion

The leech *Hirudo verbana* is an attractive system to investigate visual processing because of the animal’s known behavioral responses to complex stimuli. However, while several interneurons are known to respond to visual stimuli, their response properties are poorly understood. Among these, the S cell system is especially interesting because of its putative involvement in multimodal sensory integration (Harley et al., 2011, 2013). To improve our understanding of the role of the S cell system in visual processing, we here used a nearly intact leech preparation to quantify its spectral sensitivity under different background light conditions, to investigate spatial integration, and to test whether light adaptation is local or global.

We began by quantifying the spectral response properties of the S cell system, establishing for the first time absolute sensitivities for the leech visual system (Figure 1). We confirmed earlier reports (Kretz et al., 1976) that the leech can adapt to a wide range of background light intensities. Under each of the tested background light intensities, the response range spanned approximately 2–3 log units of stimulus light intensities (Figures 1, 3, 5, 6); fairly typical for photoreceptors across the animal kingdom (Kawamura, 1993). When fully dark adapted, leeches responded to green flashes as dim as 10^8^ photons/cm^2^/s (Figure 1), equivalent to the intensity of light on an overcast moonless night (Falchi et al., 2016).

Our physiological measurements support the existence of at least two distinct color channels (green and UV). Interestingly, the contribution of the two color channels to the response of the S cell system is dependent on the background light level, which could explain the seemingly contradictory results of previously published studies: We found that only one photoreceptor channel is active under dark background conditions and with green background illumination up to about 10^13^ photons/cm^2^/s (equivalent to twilight conditions,^1^ Figure 2). Under brighter background conditions, our results indicated that a second channel became active as well (Figures 2 and 3). This channel was predominantly UV sensitive. Both channels remained active even at the brightest green background illumination that we tested, 10^16^ photons/cm^2^/s (equivalent to full daylight). However, under this background illumination—bright green background with no UV component— the sensitivity to UV was now stronger than to green light (Figure 3). We thus both confirmed the observations of Jellies (2014a, b) and explained the apparent conflict with the earlier results of Kretz et al. (1976).

The existence of two distinct photoreceptor classes was further supported by our transcriptomic data which indicated the expression of at least two distinct opsins within the body wall of the leech (Figure 4). Similar opsins had previously been identified in another leech, *Helobdella robusta*, and comparison with opsins from other invertebrate species is compatible with these opsins being the green-sensitive and UV-sensitive receptors that underlie our physiological results. Future studies will be necessary to confirm the specific sensitivity of these specific opsins.

Many animals employ multiple photoreceptor classes that become active at different light levels; for instance in humans, rods contribute to vision most strongly at low light levels, whereas cones only become active at higher light levels (Fain and Dowling, 1973; Ingram et al., 2016). That a similar differentiation between photoreceptor classes exists in leech testifies to the complexity and richness of its sensory system.

Since the S cell system spans the entire length of the leech’s body, it appears well-positioned to integrate stimuli from different locations. We therefore investigated spatial aspects of the S cell’s responses to light. In the first series of experiments (Figure 5), we determined that adaptation to background illumination is local, suggesting that adaptation occurs in the sensory periphery or perhaps in the early stages of sensory processing. In the second series of experiments (Figure 6), we determined that the S cell system integrates stimuli from across the entire ventral body wall. We found that the maximum response of the S cell increases with the size of the illuminated area, but that the light intensity required to elicit half of that maximum response changed only marginally. This suggests that the S cell system pools (i.e., sums) responses. The existence of summation mechanisms is consistent with the organization of the S cell system, as the individual S cells are strongly coupled to each other by electrical synapses across the entire length of the body of the leech (Frank et al., 1975), so that the whole S cell system can be considered as a single syncytium that acts as a fast conducting pathway connecting the segmental ganglia (Peterson, 1984). The combination of local adaptation with global integration means that the S cell system can respond to small changes in illumination anywhere on the body, irrespective of whether that part of the body is exposed to bright background light or shade.

It has been suggested that the S cell system plays a key role in synchronizing general arousal throughout the nervous system of the leech (see Sahley et al. 1994). Related functions could potentially include an involvement in the modification or activation of motor output, and facilitating or enhancing other effects of changes in sensory input.

Taken together, our results show that the response properties of the S cell system to visual stimuli are very rich and complex, and that this system would be an ideal target for further investigations into the mechanisms and function of such complex integration.

## Acknowledgments

We wish to thank Elke Buschbeck and John Layne for the use of equipment and lab space.

## Data availability

Illumina sequencing files have been deposited to the NCBI SRA (Bioproject: PRJNA504032). Electrophysiology data and code used for analysis is available upon reasonable request.

## Supplementary information

Supplementary material, comprising one table, one figure, and two data files accompany this manuscript.

## Appendix: Converting units of light intensity

By definition, 1 W is 589 lumens at 530 nm, which is the approximate wavelength of our green light. From basic physics, 1 photon has an energy of *E* = *hc*/λ = 3.74 × 10^−19^ J. Thus a photon flux of 10^8^ photons/cm^2^/s corresponds to an energy flux of 3.74 ×10^−11^ J/cm^2^/s = 3.74 ×10^−11^ W/cm^2^ = 3.74 ×10^−7^ W/m^2^. Given 589 lumens per watt, that is equivalent to (3.74 × 10^−7^ × 589) lumens/m^2^ = 2.2 ×10^−4^ lumens/m^2^ = 2.2 ×10^−4^ lux.

## Footnotes

1 Wikipedia: https://en.wikipedia.org/wiki/Lux, retrieved January 3, 2019. For green light, 1 lux is equivalent to 4.5 ×10^11^ photons/cm^2^/s (see Appendix).

## References

Altschul SF, Madden TL, Schaffer AA, Zhang J, Zhang Z, Miller W, and Lipman DJ, 1997. Gapped BLAST and PSI-BLAST: a new generation of protein database search programs. Nucleic Acids Res 25 (17): 3389–3402. PMID 9254694.

Bagnoli P, Brunelli M, and Magni F, 1973. Afferent connections to the fast conduction pathway in the central nervous system of the leech Hirudo medicinalis. Arch Ital Biol 111 (1): 58–75. PMID 18843826.

Carlton T and McVean A, 1993. A comparison of the performance of two sensory systems in host detection and location in the medicinal leech Hirudo medicinalis. Comp Biochem Physiol Comp Physiol 104 (2): 273–277. PMID 8095878.

Desimone R and Duncan J, 1995. Neural mechanisms of selective visual attention. Annu Rev Neurosci 18: 193–222. PMID 7605061.

Dickinson MH and Lent CM, 1984. Feeding behavior of the medicinal leech, Hirudo medicinalis L. J Comp Physiol 154 (4): 449–455.

Döring C, Gosda J, Tessmar-Raible K, Hausen H, Arendt D, and Purschke G, 2013. Evo-lution of clitellate phaosomes from rhabdomeric photoreceptor cells of polychaetes—a study in the leech Helobdella robusta (Annelida, Sedentaria, Clitellata). Front Zool 10 (1): 52. PMID 24007384.

Eaton JW, Bateman D, Hauberg S, and Wehbring R, 2017. The GNU Octave version 4.2.1 manual: a high-level interactive language for numerical computations. URL http://octave.org/doc/interpreter, retrieved Jan 8, 2019.

Fain GL and Dowling JE, 1973. Intracellular recordings from single rods and cones in the mudpuppy retina. Science 180 (4091): 1178–1181. PMID 4707063.

Falchi F, Cinzano P, Duriscoe D, Kyba CC, Elvidge CD, Baugh K, Portnov BA, Rybnikova NA, and Furgoni R, 2016. The new world atlas of artificial night sky brightness. Sci Adv 2 (6): e1600377. PMID 27386582.

Frank E, Jansen JK, and Rinvik E, 1975. A multisomatic axon in the central nervous system of the leech. J Comp Neurol 159 (1): 1–13. PMID 162801.

Grabherr MG, Haas BJ, Yassour M, Levin JZ, Thompson DA, Amit I, Adiconis X, Fan L, Raychowdhury R, Zeng Q, Chen Z, Mauceli E, Hacohen N, Gnirke A, Rhind N, di Palma F, Birren BW, Nusbaum C, Lindblad-Toh K, Friedman N, and Regev A, 2011. Full-length transcriptome assembly from RNA-Seq data without a reference genome. Nat Biotechnol 29 (7): 644–652. PMID 21572440.

Harley CM, Cienfuegos J, and Wagenaar DA, 2011. Developmentally regulated multisensory integration for prey localization in the medicinal leech. J Exp Biol 214: 3801–3807. PMID 22031745.

Harley CM, Rossi M, Cienfuegos J, and Wagenaar DA, 2013. Discontinuous locomotion and prey sensing in the leech. J Exp Biol 216: 1890–1897. PMID 23785108.

Ingram NT, Sampath AP, and Fain GL, 2016. Why are rods more sensitive than cones? J Physiol 594 (19): 5415–5426. PMID 27218707.

Jellies J, 2014a. Detection and selective avoidance of near ultraviolet radiation by an aquatic annelid: the medicinal leech. J Exp Biol 217: 974–985. PMID 24265432.

Jellies J, 2014b. Which way is up? Asymmetric spectral input along the dorsal-ventral axis influences postural responses in an amphibious annelid. J Comp Physiol A 200 (11): 923–938. PMID 25152938.

Johnsen S, 2012. The optics of life: a biologist’s guide to light in nature. Princeton University Press, Princeton, NJ.

Kawamura S, 1993. Molecular aspects of photoreceptor adaptation in vertebrate retina. Int Rev Neurobiol 35: 43–86. PMID 8463064.

Kretz JR, Stent GS, and Kristan WB, 1976. Photosensory input pathways in medicinal leech. J Comp Physiol 106 (1): 1–37.

Laverack MS, 1969. Mechanoreceptors, photoreceptors and rapid conduction pathways in the leech, Hirudo medicinalis. J Exp Biol 50 (1): 129–140. PMID 5776582.

Lehmkuhl AM, Muthusamy A, and Wagenaar DA, 2018. Responses to mechanically and visually cued water waves in the nervous system of the medicinal leech. J Exp Biol 221: jeb171728. PMID 29472489.

Magni F and Pellegrino M, 1978. Patterns of activity and the effects of activation of the fast conducting system on the behaviour of unrestrained leeches. J Exp Biol 76: 123–135. PMID 712325.

Peterson EL, 1984. Photoreceptors and visual interneurons in the medicinal leech. J Neurobiol 15 (6): 413–428. PMID 6520610.

Rosendale AJ, Romick-Rosendale LE, Watanabe M, Dunlevy ME, and Benoit JB, 2016. Mechanistic underpinnings of dehydration stress in the American dog tick revealed through RNA-Seq and metabolomics. J Exp Biol 219: 1808–1819. PMID 27307540.

Sahley CL, Modney BK, Boulis NM, and Muller KJ, 1994. The S-cell—an interneuron essential for sensitization and full dishabituation of leech shortening. J Neurosci 14 (11): 6715–6721.

Schulz MH, Zerbino DR, Vingron M, and Birney E, 2012. Oases: robust de novo RNA-seq assembly across the dynamic range of expression levels. Bioinformatics 28 (8): 1086–1092. PMID 22368243.

Sievers F and Higgins DG, 2014. Clustal Omega, accurate alignment of very large numbers of sequences. Methods Mol Biol 1079: 105–116. PMID 24170397.

Simaão FA, Waterhouse RM, Ioannidis P, Kriventseva EV, and Zdobnov EM, 2015. BUSCO: assessing genome assembly and annotation completeness with single-copy orthologs. Bioinformatics 31 (19): 3210–3212. PMID 26059717.

Tamura K, Peterson D, Peterson N, Stecher G, Nei M, and Kumar S, 2011. MEGA5: molecular evolutionary genetics analysis using maximum likelihood, evolutionary distance, and maximum parsimony methods. Mol Biol Evol 28 (10): 2731–2739. PMID 21546353.

Tomina Y and Wagenaar DA, 2018. Dual-sided voltage-sensitive dye imaging of leech ganglia. Bio Protoc 8 (5): e2751. PMID 29594188.

Tsukamoto H, Chen IS, Kubo Y, and Furutani Y, 2017. A ciliary opsin in the brain of a marine annelid zooplankton is ultraviolet-sensitive, and the sensitivity is tuned by a single amino acid residue. J Biol Chem 292 (31): 12971–12980. PMID 28623234.

